# Lasting effects of a single psilocybin dose on resting-state functional connectivity in healthy individuals

**DOI:** 10.1101/2021.01.28.428377

**Authors:** Drummond E-Wen McCulloch, Martin Korsbak Madsen, Dea Siggard Stenbæk, Sara Kristiansen, Brice Ozenne, Peter Steen Jensen, Gitte Moos Knudsen, Patrick MacDonald Fisher

## Abstract

**Background:** Psilocybin is a psychedelic drug that has shown lasting positive effects on clinical symptoms and self-reported well-being following a single dose. There has been little research into the long-term effects of psilocybin on brain connectivity in humans.

**Aims:** Evaluate changes in resting-state functional connectivity (RSFC) at one-week and three-months after one psilocybin dose in 10 healthy psychedelic-naïve volunteers and explore associations between change in RSFC and related measures.

**Methods:** Participants received 0.2-0.3 mg/kg psilocybin in a controlled setting. Participants completed resting-state fMRI scans at baseline, one-week and three-months post-administration and [^11^C]Cimbi-36 PET scans at baseline and one-week. We examined changes in within-network, between-network and region-to-region RSFC. We explored associations between changes in RSFC and psilocybin-induced phenomenology as well as changes in psychological measures and neocortex serotonin 2A receptor binding.

**Results:** Psilocybin was well tolerated and produced positive changes in well-being. At one-week only, executive control network (ECN) RSFC was significantly decreased (Cohen’s d=-1.73, p_FWE_=0.010). We observed no other significant changes in RSFC at one-week or three-months, nor changes in region-to-region RSFC. Exploratory analyses indicated that decreased ECN RSFC at one-week predicted increased mindfulness at three-months (r =-0.65).

**Conclusions:** These findings in a small cohort indicate that psilocybin affects ECN function within the psychedelic “afterglow” period. Our findings implicate ECN modulation as mediating psilocybin-induced, long-lasting increases in mindfulness. Although our findings implicate a neural pathway mediating lasting psilocybin effects, it is notable that changes in neuroimaging measures at three-months, when personality changes are observed, remain to be identified.

## INTRODUCTION

Psilocybin is a prodrug of the psychedelic psilocin (4-hydroxy-N,N-dimethyltryptamine) (Nichols, 2016). Effects include profound alterations in consciousness that last approximately six hours and are characterised by perceptual alterations and synaesthesia, experiences of non-duality and transcendence, and profound changes in affect (Preller and Vollenweider, 2018). Therapeutic effects of psilocybin have been reported following between one and three moderate-to-high doses (0.025-0.42 mg/kg) in brain-related disorders including major depressive disorder (Davis et al., 2020), treatment-resistant depression (Carhart-Harris et al., 2018), obsessive-compulsive disorder (Moreno et al., 2006), terminal cancer-associated anxiety (Griffiths et al., 2016), demoralisation (Anderson et al., 2020), as well as smoking (Johnson et al., 2017) and alcohol addiction (Garcia-Romeu et al., 2019). Psilocybin is currently in phase 2b for the treatment of Treatment-Resistant Depression (COMPASS Pathways Ltd.) and in phase 2a for Major Depressive Disorder (Usona Institute).

Persistent changes in personality and mood have also been observed in healthy volunteers following a single medium-to-high dose of psilocybin. These include, for example, increases in personality traits openness and extraversion, decreases in neuroticism and increases in mindful awareness (Erritzoe et al., 2018; MacLean et al., 2011; Madsen et al., 2020). These therapeutic and personality effects appear to persist for at least months and in some cases have been reported to last more than a year (Gasser et al., 2014; Johnson et al., 2017; MacLean et al., 2011).

The medicalisation of psychedelic drugs is expanding rapidly despite a limited understanding of the neurobiology underpinning therapeutic effects. Psychological theories of psychedelic therapy such as reduced negative affect (Barrett et al., 2020), increased mindfulness (Madsen et al., 2020; Murphy-Beiner and Soar, 2020; Smigielski et al., 2019), increased cognitive flexibility (Murphy-Beiner and Soar, 2020) and reduced experiential avoidance (Zeifman et al., 2020) have been proposed, as well as increased acceptance and processing of traumatic autobiographical memories (Sloshower et al., 2020), but these have no current grounding in neurobiology. Thus, in order to maximise psilocybin’s safety and efficacy as a potential therapeutic, it is important to investigate mechanisms by which psilocybin exerts its effects.

Functional magnetic resonance imaging (fMRI) resting-state functional connectivity (RSFC) measures correlations between blood-oxygen-level-dependent signals in participants instructed to simply let their mind wander (Lee et al., 2013). Despite not being focused on any task, the brain remains organised into networks (Raichle, 2015), the character of which correlates with personality traits (Cai et al., 2020; Hsu et al., 2018) and aligns with known functional and structural topology (Straathof et al., 2019).

During the psychedelic experience, psilocybin produces a reduction in the synchronised BOLD activity of the major hubs of the default mode network (DMN) (Carhart-Harris et al., 2012; Mason et al., 2020), increases between-network RSFC (Roseman et al., 2014) and increases global RSFC across the sensory cortex while decreasing global connectivity in associative regions (Preller et al., 2020). Similarly, LSD increases RSFC between high-level association cortices, which correlates with subjective reports of ego-dissolution (Tagliazucchi et al., 2016). Although understanding the neurological basis of the acute psychedelic experience is widely informative, the long-term psychological effects of psychedelics may be distinct (Carhart-Harris et al., 2016).

Five studies to date have reported effects on human brain function after the psychoactive effects of a classical psychedelic have subsided: two studies with ayahuasca and three with psilocybin (Barrett et al., 2020; Carhart-Harris et al., 2017; Pasquini et al., 2020; Sampedro et al., 2017; Smigielski et al., 2019). Post-drug brain imaging was performed within 24 hours after the psychedelic session in all but one study (Barrett et al., 2020), during which time “afterglow” effects and potential residual drug availability confounds relating effects to lasting changes (Madsen et al., 2019; Majić et al., 2015). Barrett and colleagues reported an increase in the number of significant resting-state functional connections across the brain in 12 healthy individuals from baseline to one-week and one-month post psilocybin, hypothesising that psilocybin may increase emotional and brain plasticity. None of these previous studies evaluated correlations between change in RSFC and change in personality or other psychological traits.

In the current study we evaluated the effect of a single psilocybin dose on RSFC in 10 healthy, psychedelic naïve individuals at one-week and three months after administration, evaluating changes in within- and between-network RSFC. Further, we sought to replicate a previous finding of changes in region-to-region RSFC (Barrett et al., 2020). Lastly, in an exploratory analysis we assessed correlations between network RSFC change and variables associated with increased well-being. These included personality measures, well-being and mindfulness, which we recently showed were altered three-months after psilocybin, as well as correlated with change in neocortex 5-HT2A binding (Madsen et al., 2020). Additionally, baseline neocortex 5-HT2A binding was related to the temporal character of the psychedelic experience (Stenbæk et al., 2020). Finally, we examined whether the self-reported experience was correlated with long-term changes in brain connectivity.

## MATERIALS and METHODS

### Participants

Detailed information about participants and protocol are described in a previous study (Madsen et al., 2020) and one other study, which included these and other participants (Stenbæk et al., 2020). The study was approved by the Danish Medicines Agency (EudraCT ID: 2016-004000-61, amendments: 2017014166, 2017082837, 2018023295); and by the ethics committee for the capital region of Copenhagen (journal ID: H-16028698, with amendments). The study was preregistered at ClinicalTrials.gov (identifier: NCT03289949).

Participants were recruited from a list of individuals that expressed interest in participating in a psilocybin brain scanning study. After obtaining informed consent, participants underwent screening for somatic illness, including a medical examination, an ECG, blood screening for somatic disease, and screening for psychiatric disorders using Mini International Neuropsychiatric Interview, Danish translation version 6.0.0 (Sheehan et al., 1998). Exclusion criteria were: (1) present or previous primary psychiatric disease (DSM axis 1 or WHO ICD-10 diagnostic classifications) or in first-degree relatives; (2) present or previous neurological condition/disease, significant somatic condition/disease; (3) intake of drugs suspected to influence test results; (4) non-fluent Danish language skills; (5) vision or hearing impairment; (6) previous or present learning disability; (7) pregnancy; (8) breastfeeding; (9) magnetic resonance imaging (MRI) contraindications; (10) alcohol or drug abuse; (11) allergy to test drugs; (12) significant exposure to radiation within the past year (e.g., medical imaging investigations); (13) intake of QT-prolonging medication or electrocardiogram (ECG) results indicative of heart disease, (14) blood donation less than 3 months before project participation; (15) bodyweight less than 50 kg; (16) low plasma ferritin levels (< 12 μg/L).

### Experimental Procedures

Prior to inclusion, participants were informed about the study, including safety precautions and potential effects and side-effects of psilocybin. Before the psilocybin session, all participants met at least one of the two staff members present on the psilocybin intervention day. A urine test was used to screen for common drugs of abuse (Rapid Response, BTNX Inc., Markham, Canada) on baseline imaging days. At baseline, participants filled out questionnaires including the NEO Personality Inventory-Revised (NEO PI-R) (Costa and McCrae, 2008; Skovdahl-Hansen et al., 2004), and the Mindfulness Attention and Awareness Scale (MAAS) (Brown and Ryan, 2003; Jensen et al., 2016) and completed an MRI scan session.

On a separate day, open-label psilocybin sessions were conducted including two supporting psychologists familiar with effects of psilocybin, safety precautions and interpersonal support methods (Johnson et al., 2008). Psilocybin was administered in the morning; a number of 3 mg capsules were taken with a glass of water to approximate dose (dose: 0.2 mg/kg (n = 4) and 0.3 mg/kg (n = 6)). Participants listened to a standardised music playlist, adapted from one kindly provided by Prof. Roland Griffiths, Johns Hopkins Medicine. Music was played using a stereo system. Subjective drug intensity (SDI) was measured every 20 mins using a 0-10 Likert scale (question: ‘How intense is your experience right now? 0=’Not at all’, 10=’Very much’). Measurements were obtained from the time of drug administration to the end of the session. Participants responded orally and the supporting psychologists noted their responses. At the end of psilocybin session days, participants completed questionnaires aimed to quantify aspects of the psychedelic experience, including the 11-dimension Altered States of Consciousness questionnaire (11D-ASC) (Studerus et al., 2010) the revised Mystical Experiences Questionnaire (MEQ30) (Barrett et al., 2015), and the Ego-Dissolution Inventory (EDI) (Nour et al., 2016) (median [range]: 6.4 [5.9–7.4] hours after psilocybin intake). One-week and three-months after psilocybin administration, participants returned for MRI scan sessions identical to the baseline scan session. At three-months, participants filled out questionnaires including the NEO PI-R, MAAS, and Persisting Effects Questionnaire (PEQ) (Griffiths et al., 2006, 2011) which measures psychological changes (both positive and negative) that are subjectively perceived to be due to the psilocybin experience. Number of days between psilocybin sessions and follow-up questionnaires: mean (SD) [range] = 97.8 (11.9) [79–120 days]).

### Positron Emission Tomography (PET)

The PET data used in this analysis are the same as those reported previously (Madsen et al., 2020; Stenbæk et al., 2020). [^11^C]Cimbi-36 is an agonist radioligand selective for serotonin (5-HT) 2A (5-HT2AR) and 2C receptors (Ettrup et al., 2014). Participants completed 120-minute scans on a high-resolution research tomograph (HRRT) PET-scanner (CTI/Siemens, Knoxville, USA) at baseline and one-week following psilocybin administration. Regional time-activity curves were extracted using Pvelab (Svarer et al., 2005) from a neocortex and cerebellum region for estimation of non-displaceable binding potential (BP_ND_) using the simplified reference tissue model (Ettrup et al., 2016; Innis et al., 2007). Neocortex [11C]Cimbi-36 binding predominantly reflects 5-HT2AR binding (Finnema et al., 2014).

### Magnetic resonance imaging scan parameters

Participants completed three identical MRI scan sessions: baseline, one-week, and three-months post-psilocybin. MRI data was acquired on a 3T Prisma scanner (Siemens, Erlangen, Germany) using a 64-channel head/neck coil. A high-resolution 3D T1-weighted structural image was acquired: inversion time = 900 ms, echo time = 2.58 ms, repetition time = 1900 ms, flip angle = 9°, in-plane matrix = 256 × 256, in-plane resolution = 0.9 × 0.9 mm, 224 slices, and a slice thickness of 0.9 mm, no gap. Ten minutes of resting-state BOLD fMRI data was acquired: repetition time = 2000 ms, echo time = 30 ms, flip angle = 90°, 32 axial slices with a slice thickness of 3mm, 0.75mm gap, in-plane resolution: 3.6 x 3.6 mm, iPAT acceleration factor = 2. A gradient-echo field map of the same spatial dimensions was acquired to resolve spatial distortions due to inhomogeneities in the magnetic field (repetition time = 400 ms, echo times = 4.92 and 7.38 ms). Prior to the resting-state scan sessions, participants were instructed to close their eyes, let their mind wander freely and to not to fall asleep. Resting-state scan sessions were acquired after structural image acquisition (~15-20 minutes after scanning onset) and prior to task-related fMRI measures not described here.

### Resting-state fMRI pre-processing

Resting-state fMRI data were pre-processed using SPM12 (Penny et al., 2007). This process included slice-timing correction, realignment and unwarping, coregistration of the high-resolution T1 structural image to the fMRI data, segmentation of the high-resolution T1 structural image, applying warping parameters estimated for the high-resolution T1 into MNI space to fMRI data and smoothing with an 8mm FWHM Gaussian filter. Additional denoising of time-series data was performed in CONN (version 17.c) (Whitfield-Gabrieli and Nieto-Castanon, 2012). Time series were filtered using a bandpass filter from 0.008 to 0.09 Hz. Additionally, we performed an estimation of physiological noise sources using aCompCor: regressing out the time series (and first derivative) of the first five principal components from a decomposition of the time series from white-matter and cerebrospinal fluid voxels, separately. Additionally we regressed the time series for the six motion parameters (and first derivatives) (Behzadi et al., 2007; Whitfield-Gabrieli and Nieto-Castanon, 2012). Individual outlier volumes were identified and censored using ART (global variance threshold = 4 and composite motion threshold = 2) (http://web.mit.edu/swg/software.htm). Mean denoised time series were extracted from regions-of-interest (ROIs) for further analysis. We calculated the between-region correlation across the entire time series. Pearson’s rho correlation estimates were transformed using Fisher’s r-to-z transform (i.e., r-to-z = 0.5*(ln((1+r)/(1-r))), where *r* is the Pearson’s rho and *ln* represent taking the natural logarithm). These r-to-z values were included in statistical analyses related to connectivity strength.

### Brain atlases

Regions of interest and networks were defined using an a priori defined atlas (Raichle, 2011)□. This atlas defines 36 regions belonging to seven canonical resting-state networks: network (DMN), dorsal attention network (DAN), executive control network (ECN), salience network (SN), sensorimotor network (SMN), visual network (VN) and auditory network (AN). MNI coordinates for each network can be found in (Raichle, 2011) and in supplementary table S1. Within-network connectivity was defined as the mean connectivity between each unique pair of ROIs comprising a given network. Henceforth ECN integration and disintegration refer to increased and decreased mean within-network connectivity respectively. Between-network connectivity was defined as the mean connectivity between all ROIs from two networks where each pair of ROIs contained a region from each network. To draw comparisons with a similar previous study, we also evaluated a 268-region atlas (https://www.nitrc.org/frs/?group_id=51) described previously (Barrett et al., 2020; Shen et al., 2013).

### Statistical Analysis

All statistical analyses were calculated in R (v4.0.2) (R Studio Team, 2020). Plots were constructed using the *ggplot2* package (Wickham, 2016).

Paired t-tests were performed to investigate if there were any significant differences in ART censored volumes between timepoints (one-week vs three-months). Effects of time (one-week vs baseline or three-month vs baseline) were compared separately using paired t-tests to determine the effect of time on within- and between-network connectivity and related estimates. P-values across the 28 within- and between-network comparisons at each time point were adjusted using the Bonferroni-Holm method (Holm, 1979). We report the Cohen’s d value for each post-hoc effect evaluated.

Where an effect of psilocybin on connectivity exceeded our statistical significance threshold (p_FWE_ < 0.05), exploratory *post-hoc* analyses were performed. Due to limited statistical power stemming from a small sample, we do not draw inference on statistical significance for *post hoc* analyses, but instead report standardised effect sizes and uncorrected p-values.

#### Correlations

*Post-hoc* Pearson’s product-moment correlations were performed between change in ECN connectivity and change in MAAS, change in neocortex 5-HT2AR (i.e., [^11^C]Cimbi-36 BP_ND_) (Madsen et al., 2020) as well as measures of the acute psychedelic experience (SDI, EDI, MEQ, 11-D-ASC) and change in personality (NEO-PIR). Change in ECN RSFC was also compared with the PEQ using a linear latent variable model capturing shared covariance in individual behavioural change measures using the *lava* package (v. 1.6.8 in R (Holst and Budtz-Jørgensen, 2013)).

#### Barrett replication analysis

We attempted to replicate a previously described analysis framework applied to the Shen268 atlas (Barrett et al., 2020). As described, we applied a one-sample t-test to all ROI-to-ROI connectivity estimates and retained only those edges with a statistically significant non-zero mean connectivity after Bonferroni correction for 35778 edges tested at each time point (i.e., p_unc_ < 1.4×10^-6^). Paired t-tests evaluating change from baseline at one-week or three-months were performed for each edge surviving correction. Suprathreshold edges were identified as either increases or decreases in connectivity following psilocybin.

In our view, the type-I error for the test of interest (i.e., effect of psilocybin) is inflated by the initial “edge filtering” step considering each time point separately and further inflated by not adjusting for the family of tests of interest (695 reported in Barrett et al., 2020). Accordingly, we report paired t-tests evaluating changes from baseline to one-week or three-months, adjusting p-values using the Bonferroni-Holm method for the set of edges tested.

Code can be made available upon request.

## RESULTS

### Population

Six male and four females participated in this study (mean ± s.d. age = 28.3 ± 3.4 years). Acute psychedelic effects were well tolerated in all participants and no serious adverse events occurred. Based on self-report SDI scores throughout the sessions, the psychedelic experiences were characterized by three distinct phases, the onset, peak plateau and descent (Stenbæk et al., 2020). As previously reported, participants in this study self-reported changes in personality, including increased trait openness and mindfulness (Madsen et al., 2020). Self-reported increases in positive attributes from the PEQ (including spirituality) were 25.9% ± 21.5% (mean ± s.d.), whereas increased negative attributes reported were 1.4% ± 1.7%. Time between baseline and psilocybin intervention was 15.3 ± 9.3 days, intervention and one-week rescan was 6.5 ± 1.4 days and intervention and three-month rescan was 101.5 ± 9.9 days (mean ± s.d.).

### Lasting psilocybin effects on network connectivity

Within- and between-network RSFC structure was as expected, e.g., high within-network connectivity and relatively lower between-network connectivity (Figure 2A). Mean composite motion and censored volumes were low (3.5 ± 4.0 volumes; mean ± s.d.) and not statistically significantly different between scan times (p > 0.05).

**Figure 1.**
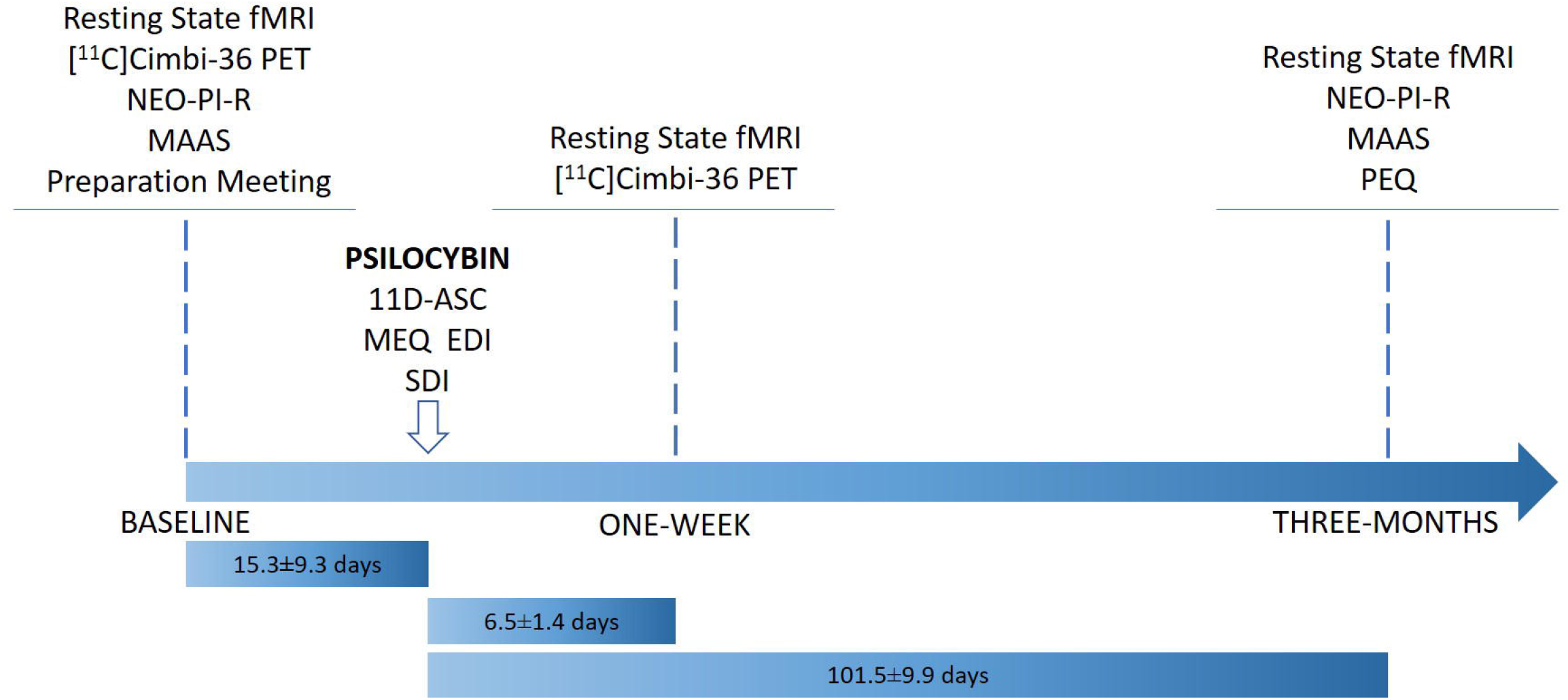
Flow chart describing study design. Note that not all data collected at a single time point was collected on a single day. Time lengths (mean ± s.d.) describe time between MRI scan sessions and psilocybin session.

**Figure 2.**
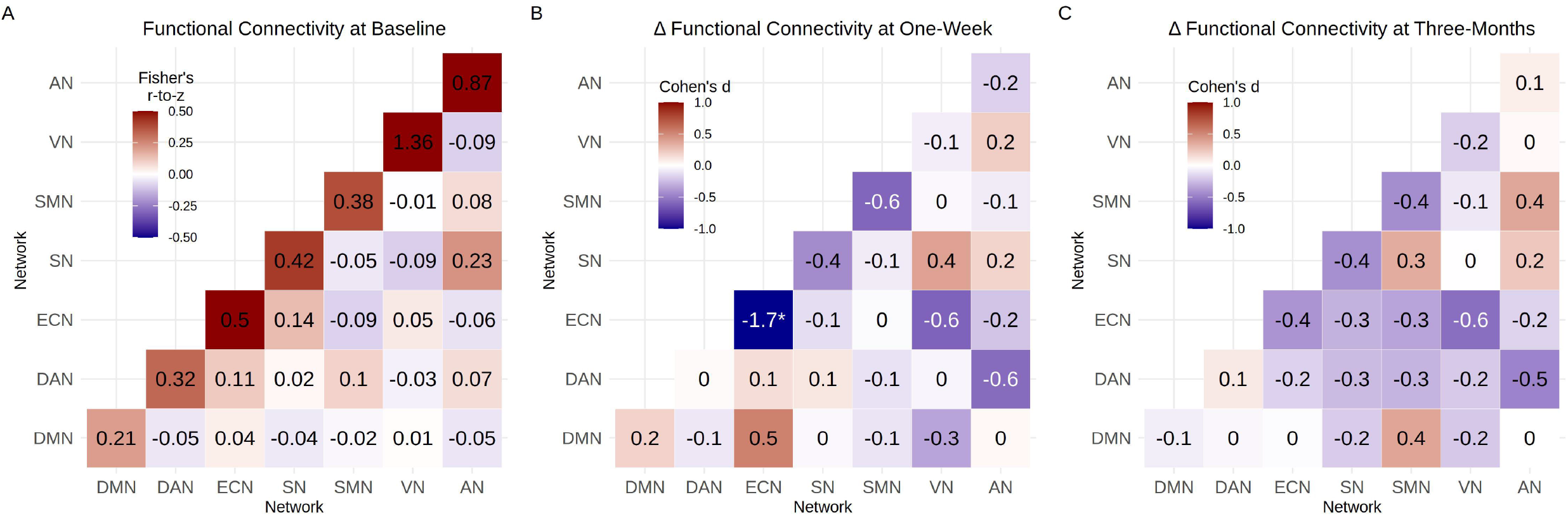
Psilocybin effects on within- and between-network resting-state connectivity. (a) Mean within- and between-network functional connectivity, cell values and colour scale represent mean r-to-z values across participants. (b) Change in connectivity from baseline to one-week rescan. (c) Change in connectivity from baseline to three-months rescan. Cell values and colour scale in (b) and (c) represent effect size (Cohen’s d). * denotes change that is statistically significant after adjustment across 28 tests (i.e., p_FWE_<0.05).

ECN within-network connectivity was statistically significantly decreased at one-week (p_unc_ = 0.00039, p_FWE_ = 0.010, Cohen’s d = −1.73; Figure 2B and 3). Nine of ten participants showed reduced ECN RSFC at one-week. At three-months, ECN RSFC remained numerically decreased as compared to baseline but this effect was not statistically significant (p_unc_ = 0.23, p_FWE_ =1, Cohen’s d = 0.4). No other within- or between-network connectivity estimates were statistically significantly altered at one-week or three-months (Table S2). At one week, SMN-SMN, ECN-VN and DAN-AN connectivity decreased (Cohen’s d = −0.6) while DMN-ECN connectivity increased (Cohen’s d = 0.5). At three-months ECN-VN and DAN-AN connectivity remained decreased (Cohen’s d = −0.6 & −0.5 respectively). Cohen’s d with magnitude > 0.5 represent a “medium” effect size (Ferguson, 2009).

**Figure 3.**
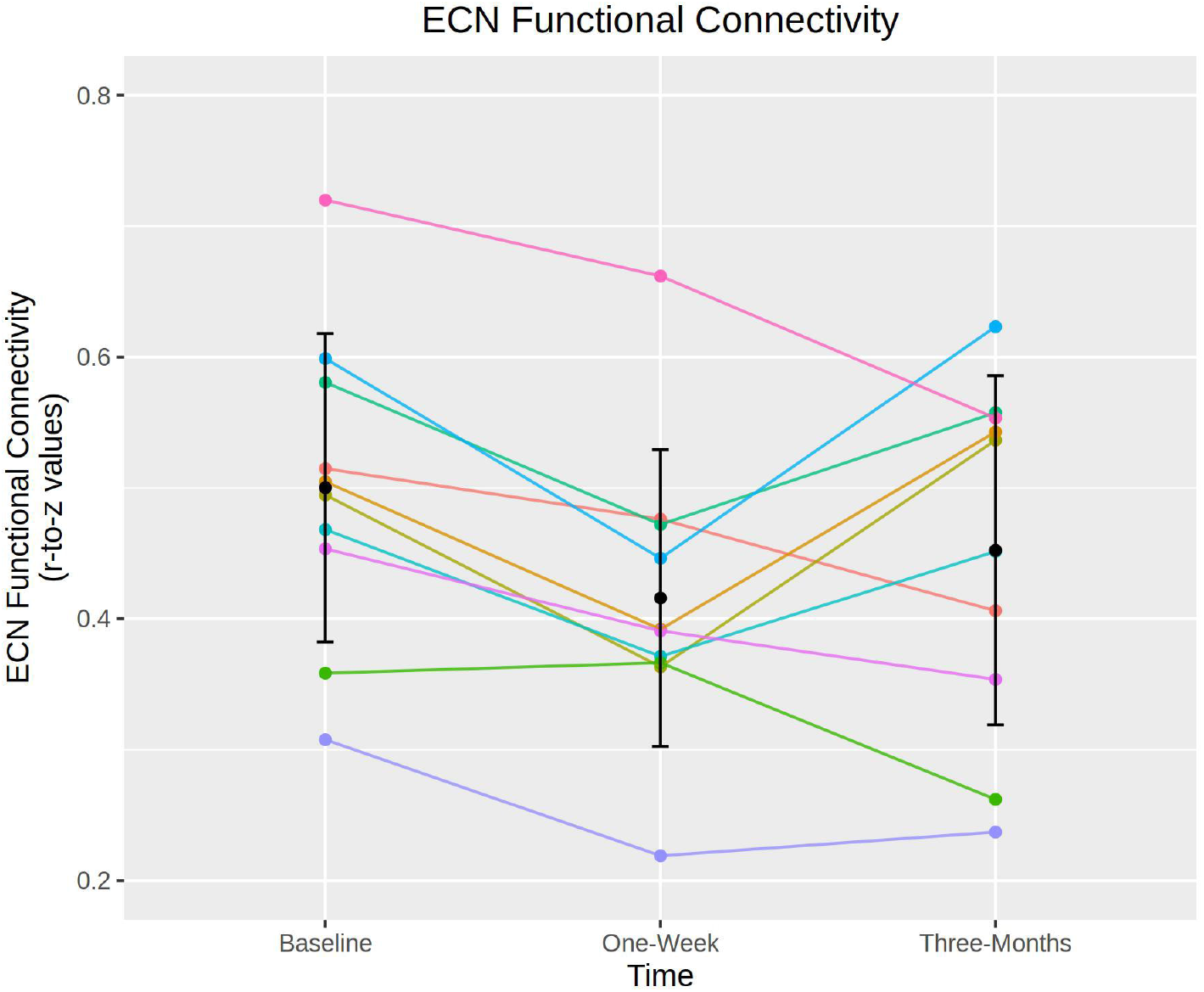
Executive control network connectivity by participant. Spaghetti plot showing individual changes in mean ECN connectivity scores (y-axis) and time point (x-axis). Error bars represent mean ± standard deviation. Colours represent individual participants. ECN connectivity is significantly decreased at one-week but not at three-months.

### Replication of previous study

Next, we attempted to replicate previously reported findings from a similar study (Barrett et al., 2020). Of the 35778 edges defined by the Shen268 atlas, 405 showed evidence for significant connectivity based on the strategy described by Barrett and colleagues, who reported 695 suprathreshold edges. RSFC was altered in 25 edges at one-week (19 increased, six decreased) and 18 edges at three-months (12 increased, six decreased) at a statistical threshold of p_unc_ < 0.05. Two of these edges were altered in the same direction at both time points (one increased, one decreased). Although we observed fewer total edges showing evidence for significant connectivity, we observed a similar proportion of edges showing a time effect (i.e., 25/405 and 18/405 are approximately similar to 48/695 and 29/695, respectively). None of the 25 nor 18 edges remained statistically significant after controlling the type-I error for the 405 tests using the Bonferroni-Holm method.

### Exploratory associations with ECN functional connectivity

Lastly, we explored the association between change in ECN RSFC at one-week and self-report measures of the psychedelic experience acquired immediately after the experience, change in neocortex 5-HT2AR (at one-week), change in personality (at three-months), and self-reported persisting effects of the psychedelic experience (three-months). A summary of correlations can be found in Table S3.

The most promising associations were observed between changes in ECN functional connectivity at one-week and three-months with self-reported change in MAAS at three-months (r [95% CI] = −0.65 [−0.91, −0.04]) and (r [95% CI] = 0.71 [0.15,0.93]) respectively, i.e., the greater the decrease in ECN connectivity at one-week, the greater the increase in MAAS score at three-months. A smaller ECN connectivity change from baseline to three-months was associated with a greater increase in MAAS score.

#### Neocortex 5-HT2AR

Change in neocortex 5-HT2AR binding measured with [^11^C]Cimbi-36 BP_ND_ at one-week was not correlated with change in ECN connectivity at one-week (r [95% CI] = 0.52 [−0.15, 0.87]). However, change in neocortex 5-HT2AR at one-week correlated more strongly with ECN RSFC change at three-months (r [95% CI] = −0.67 [−0.91, −0.06]). Put another way, greater disintegration of the ECN correlated with more neocortex 5-HT2AR.

#### Persisting Effects Questionnaire

The positive subscales of the PEQ loaded strongly onto a single latent construct, indicating high shared correlation (p < 10^-6^). Change in ECN RSFC at one-week was negatively associated with the underlying latent variable (−19.9 [−41.7, 1.93], units: change in Life Positivity PEQ per 0.1-unit change in ECN RSFC; Figure 5). Individual estimates are reported in Fig 5.

**Figure 4.**
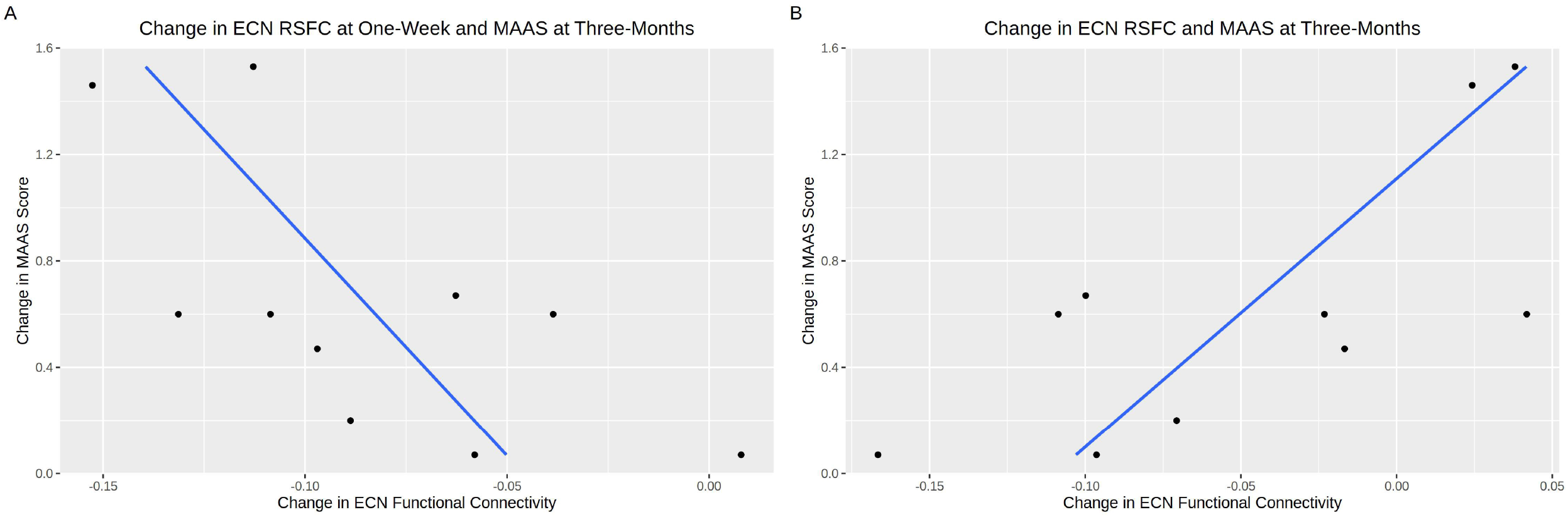
Correlations between ECN connectivity and Mindful Attention Awareness Scores (MAAS). Scatter plots with linear regressions between MAAS score (y-axis) and (a) change in ECN connectivity at one-week, and (b) change in ECN connectivity at three-months. Blue lines represent lines of best fit and black dots denote observed data.

**Figure 5.**
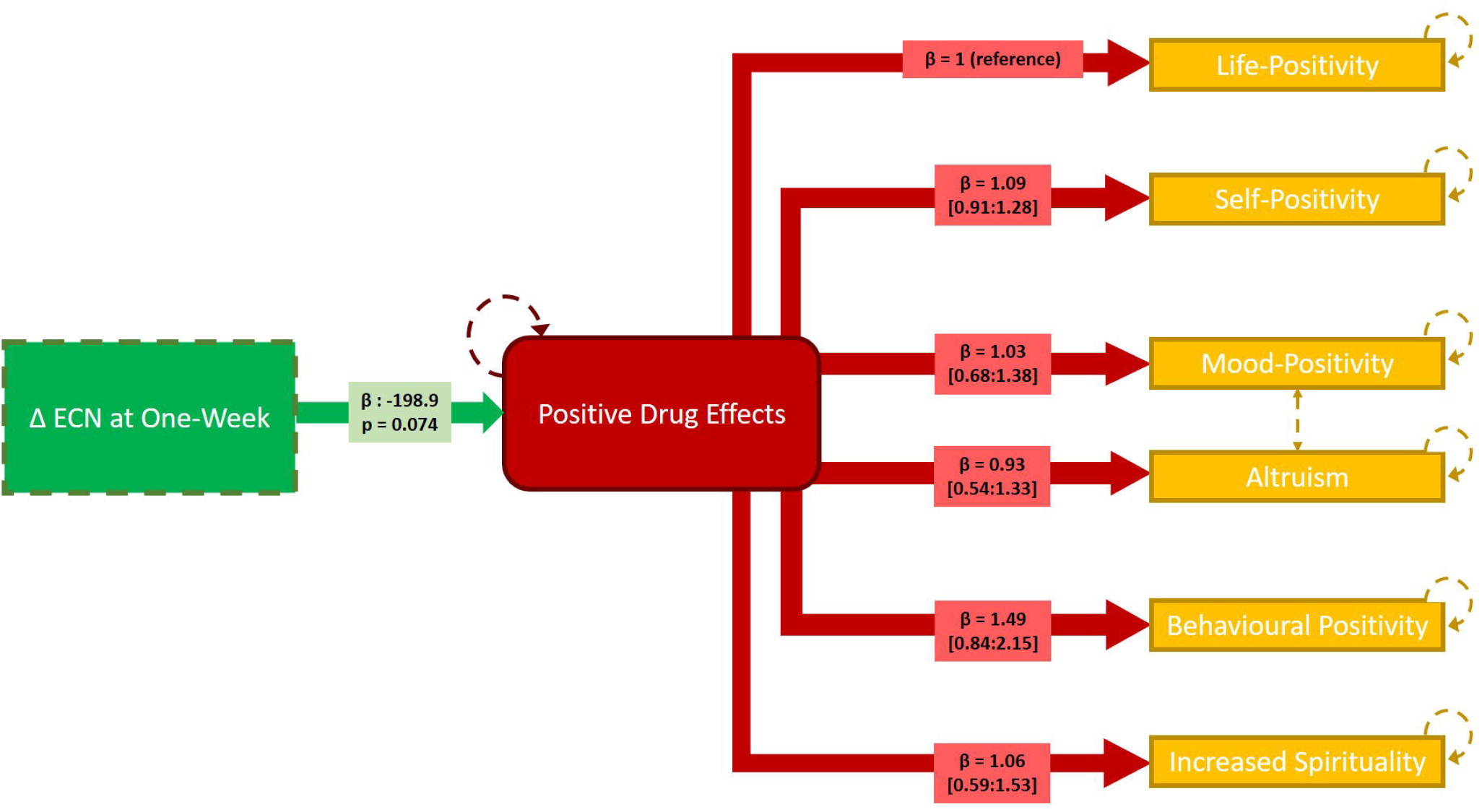
Linear latent variable model linking change in ECN connectivity at one-week and PEQ responses at three-months. The green box denotes observed change in ECN connectivity at one-week. The red box denotes the latent variable (“Positive Drug Change”). The yellow boxes denote observed PEQ scores. Hatched orange lines between “mood-positivity” and “altruism” indicate additional shared covariance. Hatched lines denote model components estimated with error. The loading parameter, β, reflecting the correlation of the score with the latent variable for each model path is noted in respective boxes (95% confidence intervals indicated for estimates between latent variable and PEQ subscale scores). Significance of the estimated effect of the ECN change on the latent variable is also noted.

## DISCUSSION

Our results show that psilocybin, when administered to healthy volunteers in a controlled environment, statistically significantly decreases ECN RSFC at one-week, but not at three-months. We observed correlations between ECN RSFC changes and changes in MAAS, neocortex 5-HT2AR and positive aspects of the PEQ, implicating alterations in ECN connectivity as a potential mechanism underlying the clinical and behavioural effects of psilocybin. No other network connectivity estimates were statistically significantly affected at one-week or three-months. Our study is small, but nevertheless implicates a candidate brain system underlying lasting psilocybin effects that can be examined in future studies in healthy and patient populations.

### Executive Control Network Connectivity

A single psilocybin administration decreased ECN RSFC; executive functions include e.g., cognitive flexibility, goal setting, attentional control and information processing (Niendam et al., 2012). This is consistent with qualitative reports in addiction patients who report persistent changes in attentional control and goal setting following psilocybin intervention (Nielson et al., 2018; Noorani et al., 2018). Though little research has been performed on ECN RSFC and its association with trait changes in executive functions, one study reported lower FC within the ECN in long-term tai-chi practitioners relative to controls. These practitioners also showed increased trait mindfulness and performed better on emotion-regulation tasks (Liu et al., 2018) aligning with our finding that decreased ECN connectivity predicted increased trait mindfulness. Change in ECN connectivity was not associated with any measures of the acute experience (Table S3) suggesting it might not simply be mediated by brain psilocin concentration (Madsen et al., 2019).

Psilocybin has been shown in small open-label trials to be extremely efficacious in the treatment of depression (Carhart-Harris et al., 2018; Davis et al., 2020), which is partly characterised by deficits in executive functions (Snyder, 2013). Our finding that psilocybin decreased ECN RSFC is consistent with a recent study reporting that unmedicated, first-time MDD patients demonstrated hyper-connectivity between the left dorsolateral PFC and frontal and parietal regions, nodes which commonly constitute cognitive control networks (Shen et al., 2015). However, another study reported that MDD is characterised by reduced frontoparietal control system connectivity (Kaiser et al., 2015). **Error! Bookmark not defined.** Although it is intriguing that in healthy individuals we observe an effect of psilocybin on a resting-state network that displays pathological connectivity in depressed patients, future studies are necessary to more clearly establish this network’s relation to treatment induced changes in measures of personality and well-being. Additionally, it remains to be established whether psilocybin induced changes in connectivity would produce the well-being states described by such connectivity signatures in healthy, untreated individuals.

The observed effect on ECN RSFC may also align with psilocybin’s potential effects on OCD (Moreno et al., 2006) and addiction (Bogenschutz et al., 2015; Johnson et al., 2017), disorders broadly characterised by aberrant control of behaviour. OCD patients display greater connectivity in ECN regions, including dorsolateral PFC (Chen et al., 2016) suggesting that reducing ECN connectivity could be therapeutically beneficial. Although individuals with addiction disorder show neuropsychological impairment in brain regions associated with cognitive control (Goldstein et al., 2004), RSFC investigations of control networks in addiction have so far utilised alternative network definitions and thus are not directly comparable to these findings (Sutherland et al., 2012).

As we only see a significant change in ECN connectivity at one-week and not at three-months, this may reflect an “afterglow” phenomenon as has been described previously (Majić et al., 2015; Murphy-Beiner and Soar, 2020). Despite the lack of long-term effects, the association between the one-week effect and long-term measures of wellbeing suggests a role for this period in mediating long-term effects on well-being, the neural correlates of which were not detected in this study. Detailed quantitative characterisation of the “afterglow” phenomenon could enable optimisation around this potentially clinically important phase of psychedelic psychotherapy by, for example, informing best practice surrounding post-session integration.

### Additional Effects on Connectivity

Besides ECN RSFC at one-week, all other standardized effect sizes for psilocybin induced change in RSFC (within- and between-network) were small to medium (i.e., |Cohen’s d| < 0.5). This suggests that future studies evaluating similar effects of psilocybin on RSFC would require sample sizes >60 to be adequately statistically powered (i.e., (1-β) > 0.8). Thus, our findings argue against large scale changes in network connectivity structure insofar as we have quantified them here. This is notable considering participants report substantive changes in personality and well-being lasting for at least one year (Erritzoe et al., 2018; MacLean et al., 2011; Madsen et al., 2020) that are likely not entirely explained by ECN changes.

Our results suggest a small effect of psilocybin on DMN RSFC (Cohen’s d = 0.19). This is noteworthy because DMN disintegration has been reported when scanning participants during the psychedelic experience (Carhart-Harris et al., 2012; Preller et al., 2020) and may be implicated in the therapeutic effects of psilocybin as DMN connectivity is elevated in a range of conditions and is reduced following administration of psilocybin to experienced meditators (Smigielski et al., 2019; Whitfield-Gabrieli and Ford, 2012). Although we do not see a persistent effect on DMN connectivity, acute disruption of the DMN may still be therapeutically relevant (Carhart-Harris and Friston, 2019; Mason et al., 2020). Intake of both Salvinorin-A, a 5-HT2AR-independent hallucinogen, and MDMA, which has a subjective effect profile distinct from serotonergic psychedelics (Doss et al., 2020; Müller et al., 2020; Roseman et al., 2014), leads to a reduction in DMN connectivity. Reduced DMN connectivity is not associated with ego-dissolution in response to psilocybin (Lebedev et al., 2015). This muted effect on DMN RSFC after the psychedelic session is consistent with a study that scanned MDD individuals one day after the psychedelic experience (Carhart-Harris et al., 2017) and healthy volunteers one-week and one-month after psilocybin (Barrett et al., 2020). Thus, convergent evidence indicates that persistent alterations in DMN RSFC are unlikely to be a critical mechanism underlying lasting psilocybin effects.

### Associations with Executive Control Network Change

Through exploratory analyses we observed three notable associations with change in ECN RSFC. Firstly, a greater decrease in ECN RSFC at one-week and a lesser decrease in ECN RSFC at three-months both correlated with increased mindfulness score at three-months. This could reflect that ECN disintegration followed by reintegration produces lasting increases in mindful awareness, a trait that is associated with reduced stress and better mood (Brown and Ryan, 2003). Secondly, building on our previous finding that change in 5-HT2AR correlated negatively with change in mindfulness (Madsen et al., 2020), we observed that a greater decrease in ECN RSFC one-week and a lesser decrease in ECN RSFC at three-months was associated with decreased 5-HT2AR at one-week. Although speculative, our findings suggest that individual change in neocortex 5-HT2AR following psilocybin administration may effect a change in mindfulness that is mediated by changes in ECN connectivity. Thirdly, decrease in ECN at one-week was positively associated with a latent construct of positive persisting effects, reflecting positive items from the PEQ, at three-months. This effect was particularly pronounced regarding “behavioural positivity”, aligning with qualitative reports from patients in clinical trials (Nielson et al., 2018; Noorani et al., 2018). Although exploratory, these observations provide a framework for linking brain and behavioural changes effected by psilocybin administration in future studies in healthy and clinical cohorts. Although we see changes in personality-trait openness, neuroticism and conscientiousness in this sample (Madsen et al., 2020), these changes were not correlated with change in ECN RSFC (Table S3).

### Barrett Replication

Replication is critical for identifying reliable brain markers of psilocybin effects. Here we sought to replicate recently reported findings from a study very similar to ours (Barrett et al., 2020). We replicated the scale of region-to-region RSFC estimates that were affected by psilocybin. However, none of these effects remained statistically significant when controlling for multiple comparisons. Ours and the previous study have small sample sizes which exacerbate the statistical power limitations of an exploratory region-to-region RSFC analysis strategy. Without substantively larger samples, this analysis framework would likely benefit from a hypothesis-driven evaluation of specific region pairs.

### Limitations

As noted previously, our sample size of 10 individuals limits statistical power. Nevertheless, the data reported here provide a firmer foundation for future studies in clinical and healthy cohorts with larger samples. Although psilocybin seems to have positive behavioural and mood effects in healthy individuals, it is not clear how closely our observed effects on brain connectivity would generalize to clinical cohorts. Additional studies in patient groups are needed to delineate the neurobiological basis of therapeutic effects of serotonergic psychedelics including psilocybin. There is variation across resting-state atlases in how cognitive control networks are defined, limiting our ability to draw firm conclusions with previous related studies (Raichle, 2015; Schaefer et al., 2018; Shen et al., 2020). Alternative analytic strategies (e.g., dynamic functional connectivity, entropy analyses) or task-based fMRI may reveal more pronounced effects on brain function and connectivity than those reported here. Future studies integrating fMRI with PET markers may offer deeper insights into the neurobiological mechanisms mediating psilocybin effects on behaviour (Fisher and Hariri, 2012). Although the observed effect on ECN may be related to individual differences in pharmacodynamics (i.e., drug availability and metabolism), we did not measure individual plasma psilocin levels (Madsen et al., 2019).

In summary, we report effects of a single psilocybin administration on resting-state functional connectivity networks at one-week and three-months in a cohort of 10 individuals. Although a small sample, we identified a statistically significant reduction in executive control network RSFC at one-week but not at three-months (although numerically decreased). Exploratory correlations with change in ECN at one-week suggest it may be associated with change in neocortex 5-HT2AR at one-week as well as change in mindfulness and persistent positive psychological effects at three-months. Nevertheless, future studies are necessary to more thoroughly map psilocybin effects on to changes in brain function and connectivity that may mediate its lasting clinical and behavioural effects.

