## Supplementary material for "Lasting effects of a single psilocybin dose on resting-state functional connectivity in healthy individuals": Figure S1, Table S1, Table S2, Table S3

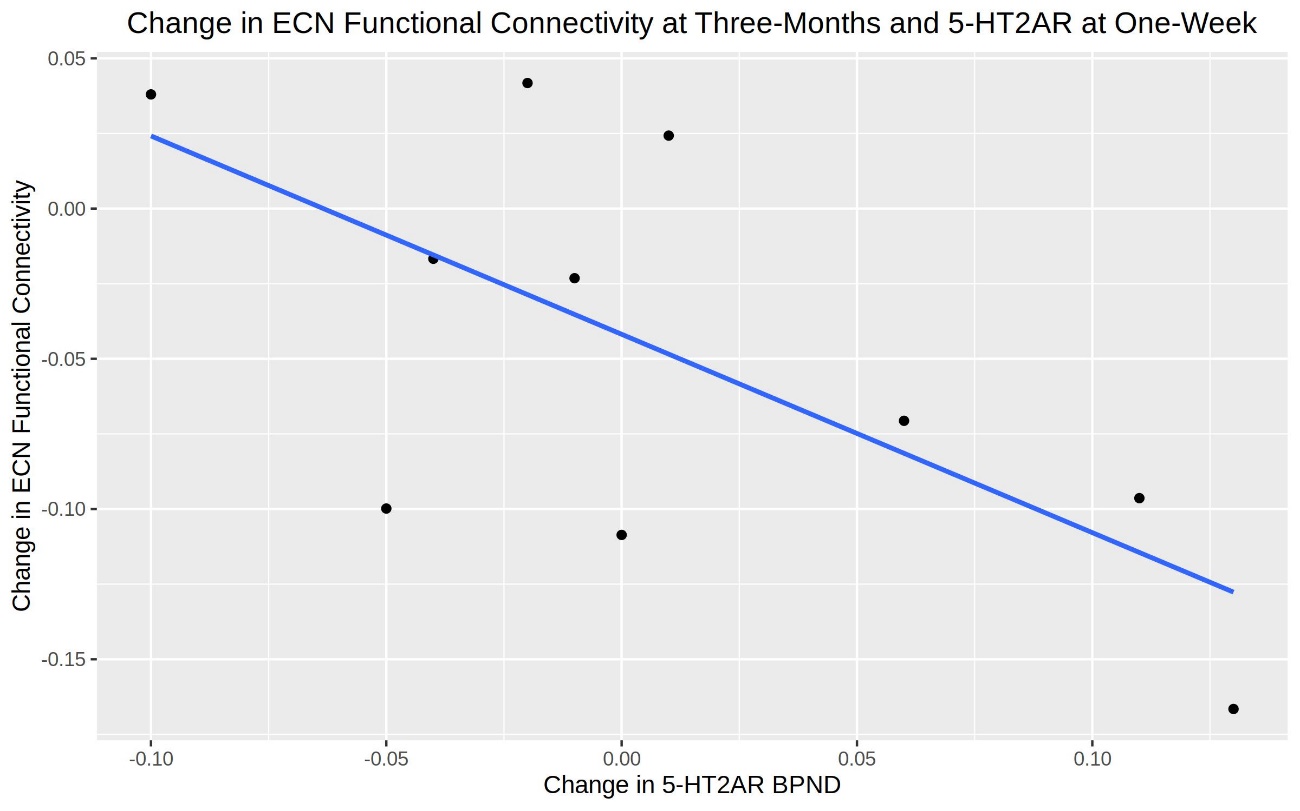

Figure S1. Correlation between the change in neocortex [^11^C]Cimbi-36 BPND and change in ECN connectivity. Scatter plot showing the change in [^11^C]Cimbi-36 BP_ND_ at one-week (x-axis) and change in ECN connectivity at three-months (y-axis). The blue line represents the line of best fit and black dots denote observed data.

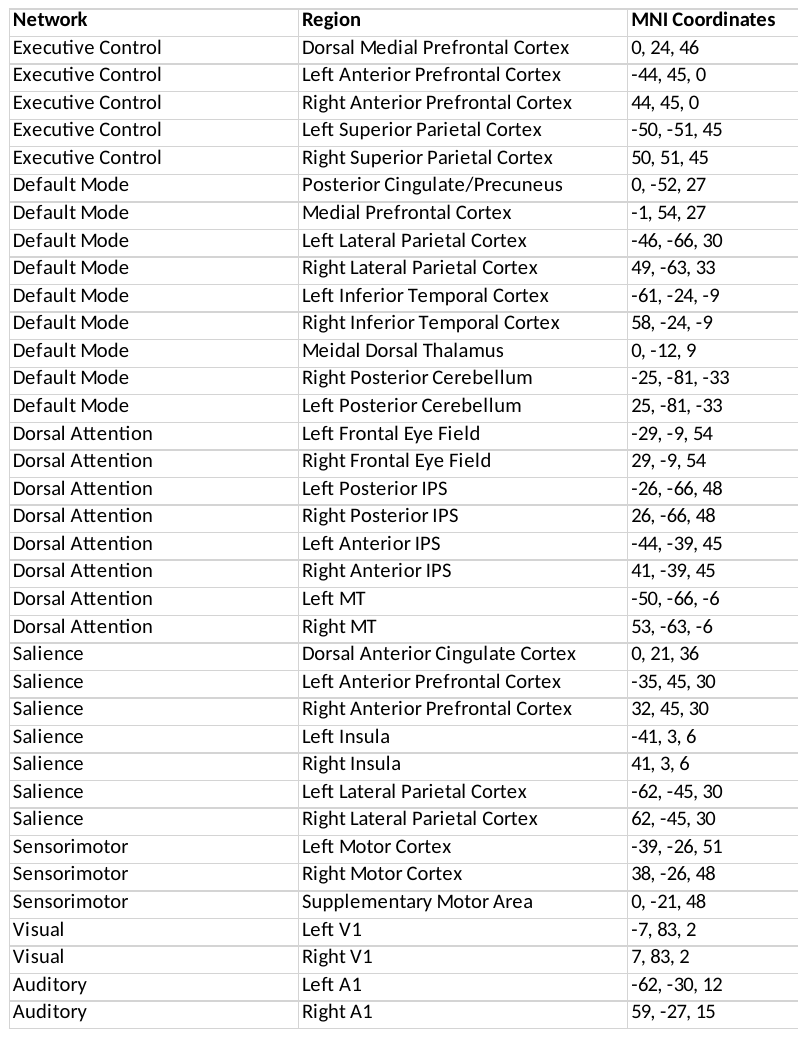

Table S1. Regions and associated brain networks as described in (Raichle, 2011).
ROIs described by 10 mm radius spheres centred on given MNI coordinates.

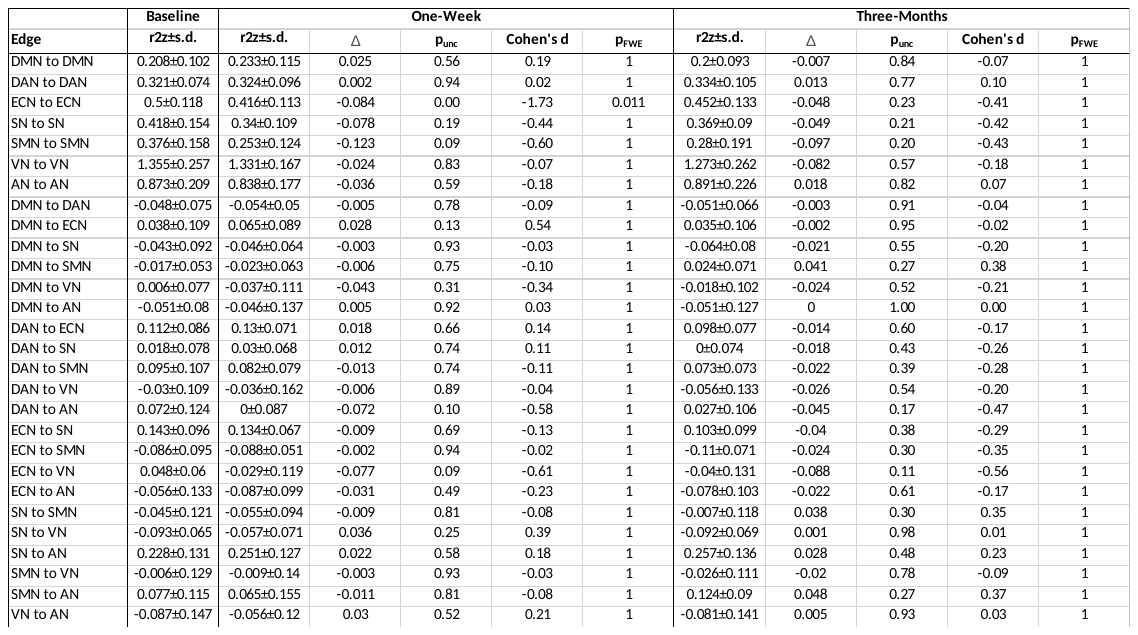
Table S2: Functional connectivity scores. DMN, Default Mode Network; DAN, Dorsal Attention Network; ECN, Executive Control Network; SN, Salience Network; SMN, Sensorimotor Network; VN, Visual Network; AN, Auditory Network.

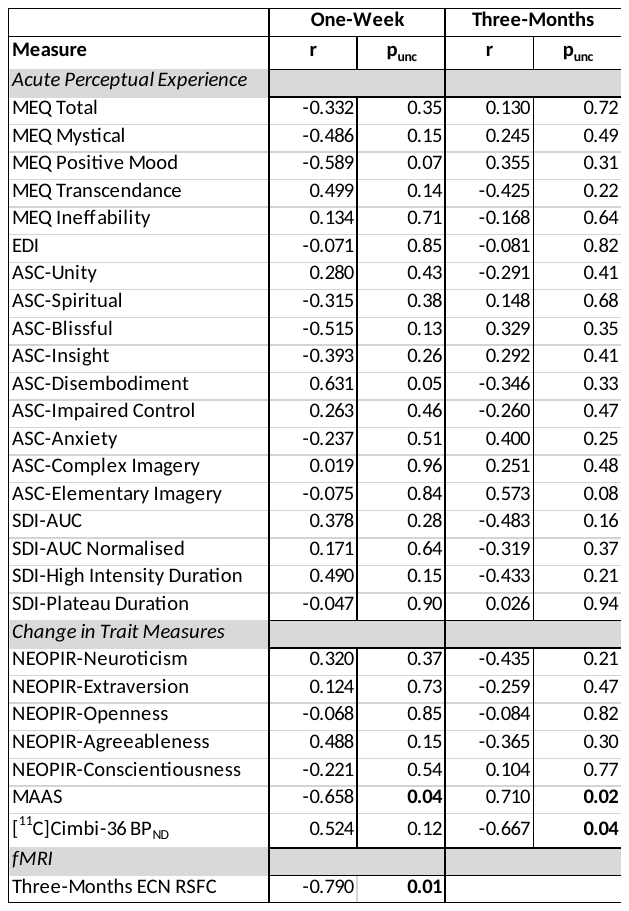

Table S3: ECN Correlations. Bold numbers highlight p_unc_ < 0.05. MEQ, Mystical Experience Questionnaire; EDI, Ego-Dissolution Inventory; ASC, Eleven Dimensions of Altered States of Consciousness Rating Scale; SDI, Subjective Drug Intensity; NEOPIR, NEO Personality Inventory Revised; MAAS, Mindful Attention Awareness Scale; BPND, Non-Displaceable Binding Potential; LCI, Lower Confidence Interval; UCI, Upper Confidence Interval; ECN, Executive Control Network; RSFC, Resting State Functional Connectivity.
